# CD8 T lymphocytes redeploy embryonic cell cycle control mechanisms to facilitate rapid cell proliferation

**DOI:** 10.1101/2025.02.28.640744

**Authors:** D. A. Lewis, A. Kar, A. Savage, V. Kelly, D. Wright, R. Zamoyska, T. Ly

## Abstract

Rapid proliferation and expansion of cytotoxic CD8 T lymphocytes is crucial for adaptive immunity against viral infection. CD8 T cell cycles complete cell division cycles in <6 hours, representing a physiological extreme for somatic mammalian cells. Embryonic stem cells also rapidly divide and have been shown to utilize specialized cell cycle control mechanisms that differ from somatic fibroblasts, including subdued periodicity of cyclin proteins. CD8 T cell cycle control remains poorly understood compared to embryonic and other somatic cell types. Here, we tested whether CD8 T cells utilize similar control mechanisms to embryonic stem cells to promote rapid cell cycles. We used mass spectrometry-based proteomics to comprehensively measure protein abundances in G1, S and G2&M phases somatic mouse CD8 T cells, mouse embryonic stem cells (mESC) and mouse fibroblasts (NIH3T3). We isolated cell cycle phases using PRIMMUS, thereby avoiding potential artefacts due to arrest-based synchronization. We discovered striking similarities between mESC and CD8 T cells. Similar to mESC, Cyclin E1 was expressed at high levels and constitutively across the cell cycle in CD8 T cells. Overall, cyclins and cell cycle regulated proteins were present in higher abundance in CD8 T cells and mESC as compared with NIH3T3 cells. Additionally, CD8 T cells express high levels of Emi1/Fbxo5 to promote S-phase entry. Interestingly, Emi1 deletion unexpectedly resulted in changes in markers of CD8 T cell phenotype, suggesting an association between cell cycle control and immune function. Thus, we show several adaptations of cell cycle control of embryonic stem cells are redeployed in somatic T cells to promote rapid cell cycles.

## Introduction

The structure and control of cell division cycles can differ between cell types in multicellular organisms to coordinate with developmental stage and the function of the cell type^1–8^. T lymphocytes that express the cell differentiation marker 8 (CD8) receptor on their cell surface (“CD8 T cells”) are specialized cells of the adaptive immune system^9^. CD8 T cells are specialized killer cells that target virally infected cells and tumour cells for elimination as part of the adaptive immune response. CD8 together with the T cell receptor (TCR) initiates T cell activation. This activation is antigen specific. CD8 binds to the constant α3 domain of MHC class I molecules and the TCR recognizes antigenic peptides bound by MHC class I. An adult human will have millions of clones of CD8 T lymphocytes, each equipped with a unique TCR to recognize pathogen-associated antigens. The frequency of each clone is estimated to be very low: ∼10^-5^ in mouse^10^ and ∼10^-6^-10^-4^ in human^11^. Following antigen recognition and activation, quiescent, naïve CD8 T cells rapidly proliferate to expand the antigen-specific clonal pool^12,13^. Most activated cells then differentiate into effector CD8 T cells that mediate cytotoxic killing, with a minority become long-lived memory CD8 T cells that are important in adaptive immunity^14^. In parallel, viral replication will result in increases in pathogen burden. Therefore, timely and rapid proliferation is crucial to expand CD8 T cell numbers and combat disease.

Like activated CD8 T cells, embryonic stem cells (ESCs) are multipotent and capable of rapid proliferation. ESC cell divisions are characterized by a specialized cell cycle control network^1,3^ that produces an altered expression profile for cell cycle promoting genes, a truncated G1 phase^6,15^, and rapid S-phase entry. In mouse embryonic fibroblasts, Cyclin E, Cyclin A, and other targets of the E2F transcription factors exhibit periodic expression across the cell cycle peaking in abundance at the G1/S transition. Cyclins E/A in complex with cyclin-dependent kinase 2 (CDK2) then hyperphosphorylates the tumour suppressor retinoblastoma protein (Rb) to relieve repression of E2F and thereby promote cell cycle gene expression^16^. In contrast, in mouse ESCs (mESCs), Cyclins E/A show dampened periodicity or constitutive expression across the cell cycle^3,4,17–19^, leading to constitutive hyperphosphorylation of Rb and rapid S-phase entry. Mechanistically, the levels of Cyclins E and A are sustained in mESCs by inhibition of the key E3 ligase, the APC/C, by its protein inhibitor, Emi1 (*Fbxo5*)^20,21^. APC/C targets hundreds of proteins for proteasomal degradation^22,23^, including the E2F targets Cyclin A, and Skp2, which mediates degradation of Cyclin E. Emi1 is itself an E2F target. Emi1 switches from being an APC/C substrate to an APC/C pseudo substrate inhibitor in a manner dependent on Emi1 protein levels^24,25^.

Given the importance of rapid clonal expansion in adaptive immunity, studies investigating cell cycle control in CD8 T cells are comparatively limited^26^. Genetic evidence points to specific adaptations in CD8 T cell cycle control. For example, in contrast to the established model that canonical E2F genes (E2F1/2/3) are pro-proliferative, E2f1/E2f2 double knockout CD8 T cells show hyperproliferation^27^. Emerging evidence suggests interplay between CD8 T cell cycle control and immune function. CD62L, a cell surface protein, is used as a marker for CD8 T differentiation, being low in effector (CD62L^lo^) and high in memory-like (CD62L^hi^) CD8 T cells. CD62L expression is highly correlated with proliferative state in activated CD8 T cells during an anti-viral response *in vivo*^12^. Following activation, CD8 T cells rapidly divide and show concomitantly decreased CD62L expression. However, over time, a subpopulation of cells divides more slowly and shows high CD62L expression^12^. CDK4/6 phosphorylates NFAT to suppress interferon gamma expression in CD8 T cells^28^. CDK4/6 inhibition also delays CD8 T cell cycle progression, enhances CD62L expression, and promotes T cell memory^29,30^. It remains unclear if this pro-memory effect is related to the known functions of CDK4/6 in promoting G1 progression by phosphorylating Rb^29^, or if this reflects a new function of CDK4/6, for example, by regulating Myc activity^30^. Together these studies, and others^31,32^, show strong correlative and causal relationships between cell cycle control and immune cell phenotypes that warrant further mechanistic investigation into rapid CD8 T cell cycles.

## Results

### CD8 T cells and mESCs share similar doubling rate and cell cycle phase frequencies

We first compared murine CD8 T cell proliferation with mouse embryonic stem cells (mESCs) and mouse NIH3T3 fibroblasts. We also included in the comparison immortalized retinal pigment epithelial cells (RPE1), which is a well characterized human cell line model used to study cell cycle control. Between 24 h and 48 h following activation, CD8 T lymphocytes underwent up to five cell divisions (Fig. 1a) and had an average cell doubling time of 7.5 hrs (Fig. 1b). The average cell doubling time of mESCs was 10 hrs (Fig. 1b). Both CD8 T lymphocytes and mESCs had significantly shorter doubling times than RPE1 (15 hrs, Fig. 1b). While differences in cell cycle timing between species exist^33^, within a species there are significant differences in the average cell doubling time of embryonic stem cells and differentiated cell types. For example, mouse embryonic fibroblasts (MEFs) have a cell doubling time of ∼48 hours^34^. Indeed, the NIH3T3 MEF cell line showed a doubling time of 22.5 hours (Fig. 1b).

**Figure 1:**
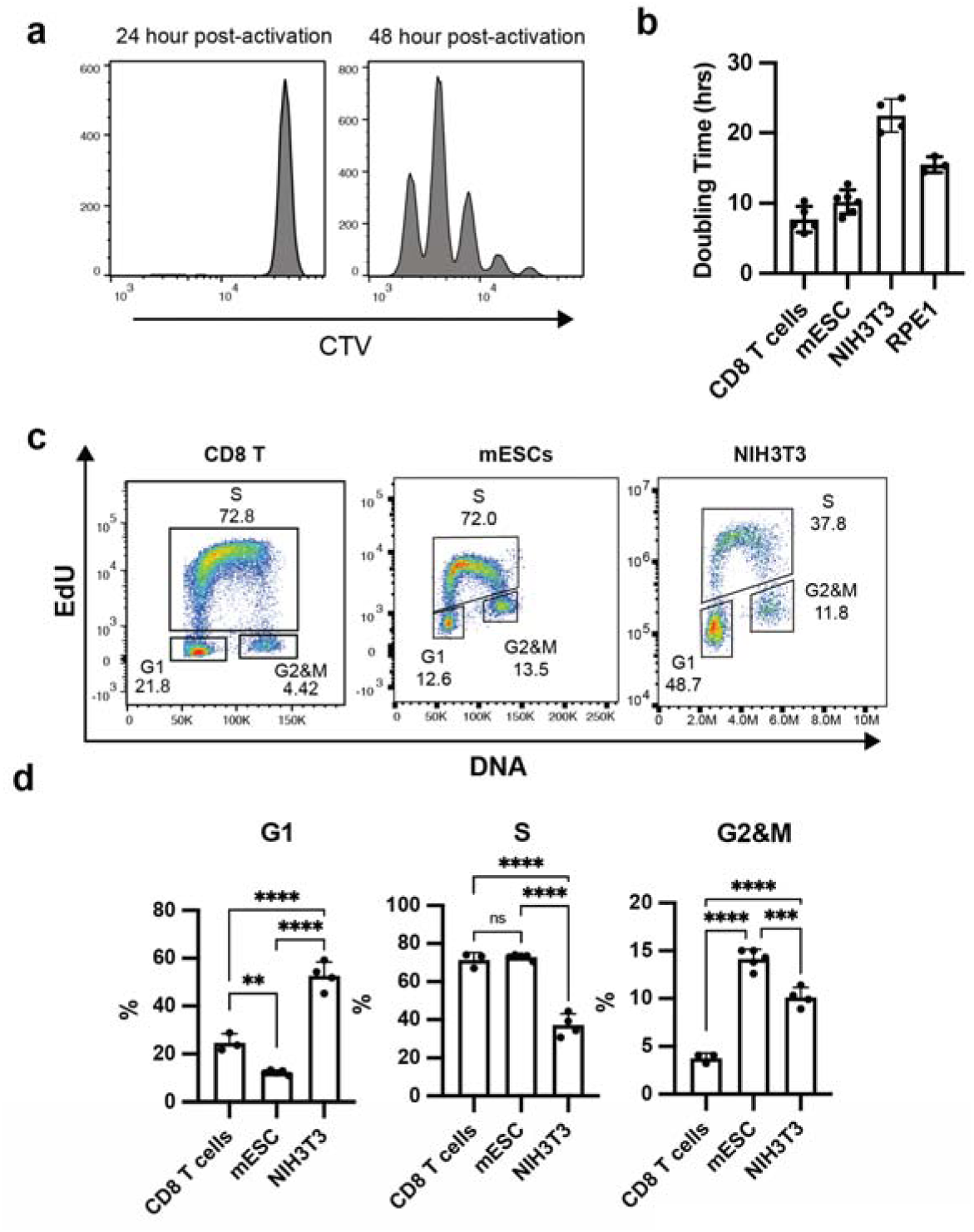
CD8 T cells are highly proliferative and are enriched in S-phase cells. **a** Naïve, quiescent CD8 T cells were stained with CellTraceViolet (CTV) prior to activation and were analyzed at indicated times post activation. **b** Doubling time was estimated from daily cell counts for CD8 T cells, mouse embryonic stem cells (mESCs), NIH3T3 fibroblasts and RPE1 cells. Results are from 3-6 biological replicates conducted over 2 independent experiments. **c** Asynchronous CD8 T cells, mESCs, and NIH3T3 cultured in exponential growth regime were pulsed with EdU for 30 minutes, harvested and subjected to Click Chemistry-based fluorescence detection. **d** Percentage of G1, S and G2&M phase cells (n = 3). Statistical significance was determined by unpaired Student’s t-test, ** <0.01, ***<0.001, ****<0.0001

The difference in average cell doubling time could reflect a difference in time spent in G1. Discrimination by DNA content and EdU labeling showed a marked increase in S-phase cells in CD8 T lymphocytes and mESCs compared to NIH3T3 (Fig. 1c, d). This increase was primarily in early S-phase cells, which have 2N DNA content and are EdU+ (Fig. 1c). CD8 T cells showed a markedly lower frequency of cells in G2&M phase (Fig. 1d). These results demonstrated that mESC and CD8 T are comparable in doubling times, with a cell cycle characterized by a shortened G1 phase and differences in G2&M duration.

### Cell type plasticity in cell cycle regulated protein abundance

Having established that the CD8 T cell cycle phase distribution resembled mESCs, we next investigated if the CD8 T cell cycle control network is convergently configured to support a truncated G1 phase and rapid cell divisions. It was previously shown that there is diminished periodicity of cell cycle regulated (CCReg) proteins in mESCs to support rapid cell divisions. These proteins included key regulators such as Cyclin A2. However, conflicting results have been obtained previously, varying from no changes^4,5,8^ to muted changes in CCReg protein abundances^3^, as recently reviewed^1^. These discrepancies have been ascribed to challenges in obtaining high synchrony in mESCs using conventional drug-based arrest and release approaches and the narrow linear range for immunoblot protein quantitation^3^.

To re-evaluate potential cell type plasticity in cell cycle regulation, and to test if CD8 T cells rely on similar adaptions to mESCs, we performed quantitative mass spectrometry-based proteomics to measure protein abundances across an unperturbed cell cycle in mouse CD8 T lymphocytes, mESC, and NIH3T3 cells. We utilized PRoteomic analysis of Intracellular iMMUnolabelled cell Subsets (PRIMMUS)^35,36^. In PRIMMUS, asynchronous cells in exponential growth are fixed in formaldehyde and immunostained for cell cycle phase specific markers, with each cell cycle phase separately isolated by FACS for downstream proteomics sample preparation. Thus, PRIMMUS avoids potential artefacts due to cellular stresses that caused by artificial synchronization as opposed to cell cycle per se^37^ and moreover, cell cycle synchronization protocols are not established for CD8 T lymphocytes. Importantly, synchronization approaches can cause differentiation in embryonic stem cells due to crosstalk between cell division and cell differentiation pathways^38^ that are avoided by using PRIMMUS.

Fixed cells were sorted into G1, S and G2&M populations, subjected to in-cell digestion^36^ and data-independent acquisition (DIA) LC-MS analysis (Fig. 2a and Methods). 8,178 protein groups were identified with 2 or more peptides per protein. Replicates were highly correlated (Pearson’s r > 0.8), and correlation within a cell line was higher than between cell lines (Supplementary Table 1). Protein intensities were normalized by dividing the intensity of each protein by the summed total protein intensities per sample and multiplying by a million to generate normalized protein abundances in parts-per-million (ppm) units (Supplementary Table 2).

**Figure 2:**
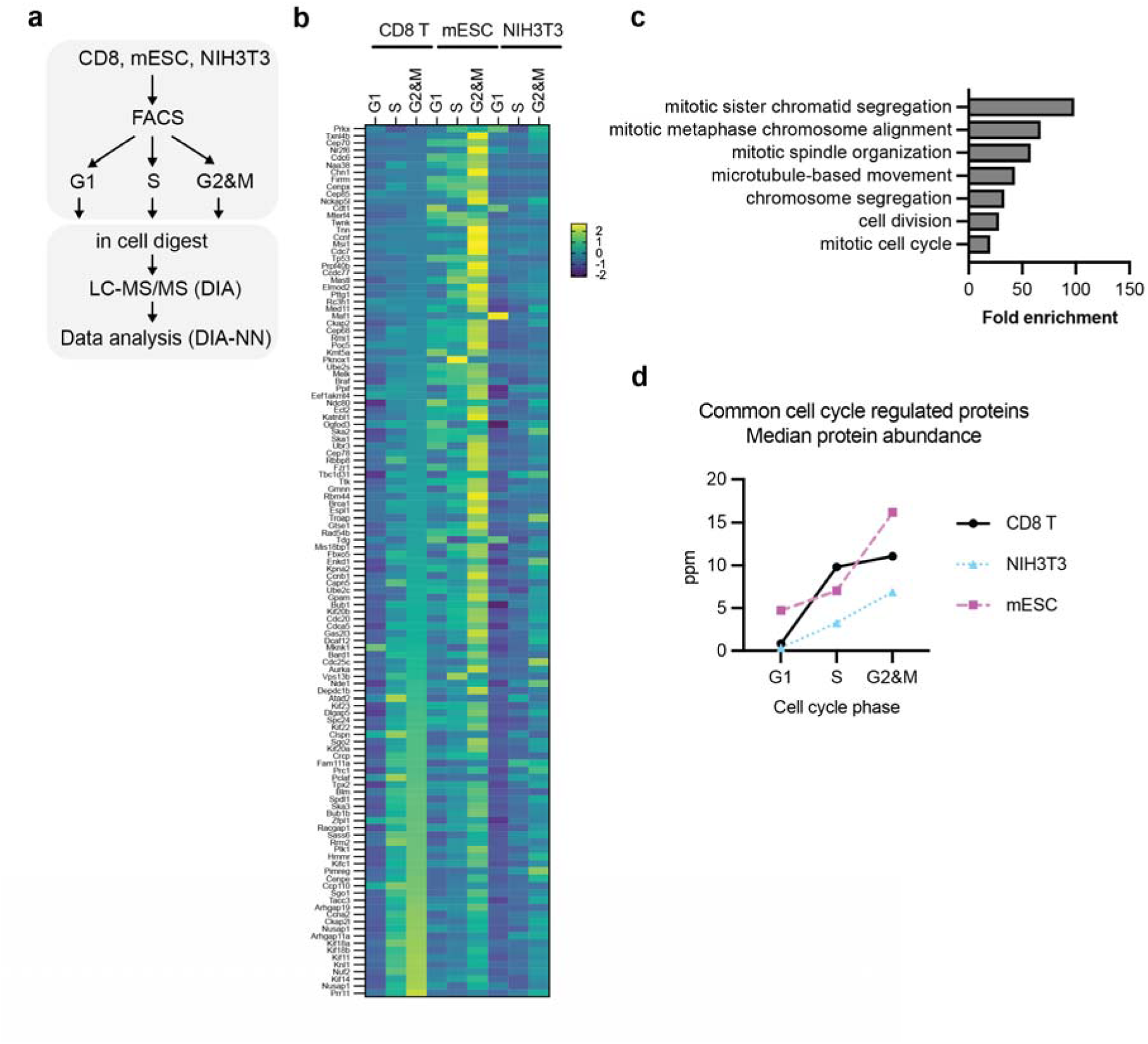
The common cell cycle regulated protein network in CD8 T, mESC and NIH3T3 cells. **a** Design of mass spectrometry-based proteomics experiment. **b** Heatmap showing the scaled protein abundances for the 121 proteins that are cell cycle regulated in CD8 T cells, mESC and NIH3T3. **c** GO term enrichment analysis of the common cell cycle regulated proteins. All GO terms shown are significantly enriched (FDR < 0.05). **d** Median abundances of the common cell cycle regulated proteins across the cell cycle phases.

First, we systematically examined proteins that were cell cycle regulated. There was a core set of 121 proteins that were consistently cell cycle regulated across all three cell types (Supplementary Table 3, Fig. 2b). These proteins were highly enriched in annotated cell cycle functions (Fig. 2c), validating the PRIMMUS strategy. In general, cell cycle regulated proteins were more abundant in the faster dividing cells in G1 and S phases of the cell cycle (Fig. 2d). In mESC, there was higher abundance in G1 phase compared to CD8 T cells and NIH3T3 (Fig. 2d). We conclude that cell cycle regulated proteins in mESC are highly periodic, and indeed, fluctuated to a similar degree compared to slower dividing NIH3T3 cells. However, significantly higher levels of these proteins were detected in G1 phase, consistent with previous reports of reduced APC/C activity in mESCs^3^.

Next, we examined whether key proteins, including cyclins, were constitutively or periodically expressed in these cell lines. Cyclin A2 showed cell cycle regulated abundance in all three cell lines (Fig. 3a). However, the abundances were generally higher in CD8 T cells and mESCs compared to NIH3T3. In NIH3T3 cells, Cyclin E1 levels were consistently detected only in S phase (Fig. 3b), consistent with a role of Cyclin E1 in promoting G1/S transition. In contrast, in mESCs and CD8 T cells, Cyclin E1 protein was robustly detected in G1, S, and G2&M phases, and at levels higher than maximally observed in NIH3T3 (Fig. 3b). Thus, while most proteins showed periodic fluctuation in all three cell lines, Cyclin E1 showed constitutive expression only in faster dividing cells suggesting Cyclin E1 as a candidate driver of rapid cell divisions.

**Figure 3:**
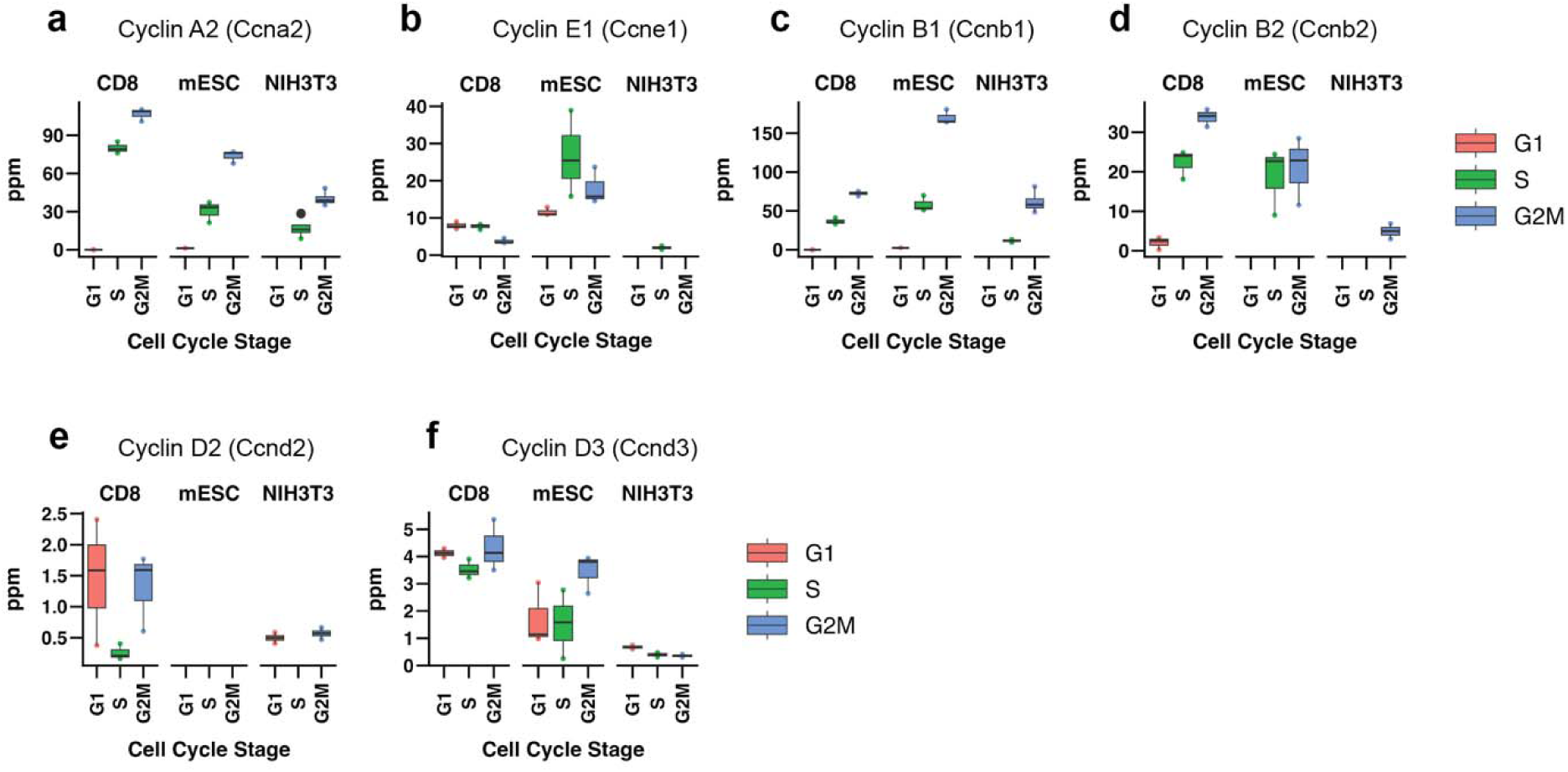
Comparing cell cycle regulation of Cyclin proteins in fast and slow dividing cell types. **a-f** Protein intensities (ppm) for Cyclin A2 (a), Cyclin E1 (b), Cyclin B1 (c), Cyclin B2 (d), Cyclin D2 (e) and Cyclin D3 (f). The middle line shown within boxplots indicate mean values (n = 3-4).

Cyclin B1 and Cyclin B2 were cell cycle regulated in all three cell lines (Fig. 3c-d) and showed maximal abundance in G2&M. Cyclin B2 levels were lower in NIH3T3 cells compared to CD8 T cells and mESCs (Fig. 2d). Cyclin B1 and Cyclin B2 show differential localization in HeLa cells, with Cyclin B2 localized to the Golgi apparatus and centrioles^39,40^. However, there is functional redundancy between the two B-type Cyclins^41^. These results would be consistent with higher overall Cyclin B activity in faster dividing cells, or specialized functions of Cyclin B2, to promote CDK activity in Golgi and centrosomal compartments in CD8 T cells and mESC.

Cyclin D2 and Cyclin D3 showed cell type enhanced expression (Fig. 3e-f). Cyclin D2 was expressed in CD8 T cells and NIH3T3 (Fig. 3e). Cyclin D3 levels were higher in CD8 T cells and mESCs (Fig. 3f). D-type cyclins in general are higher in abundance in CD8 T cells (Fig. 3e-f). The enhanced expression of Cyclin D3 in CD8 T cells is consistent with genetic studies showing tissue-specific defects in the development of the hematological system in the absence of all D-type Cyclins and in T lymphocytes in Cyclin D3 knockout mice^42,43^.

Cyclins form active kinase complexes with cognate CDK proteins. CDK abundances typically do not show the same degree of periodic expression as Cyclin proteins. Consistent with this idea, our proteomics analysis showed relatively little to no variation in CDK protein abundance across the cell cycle in these three cell types with one exception (Fig. 4a-e). Cdk4 showed higher levels in G2&M compared to G1 phase in mESC (Fig. 4a).

**Figure 4:**
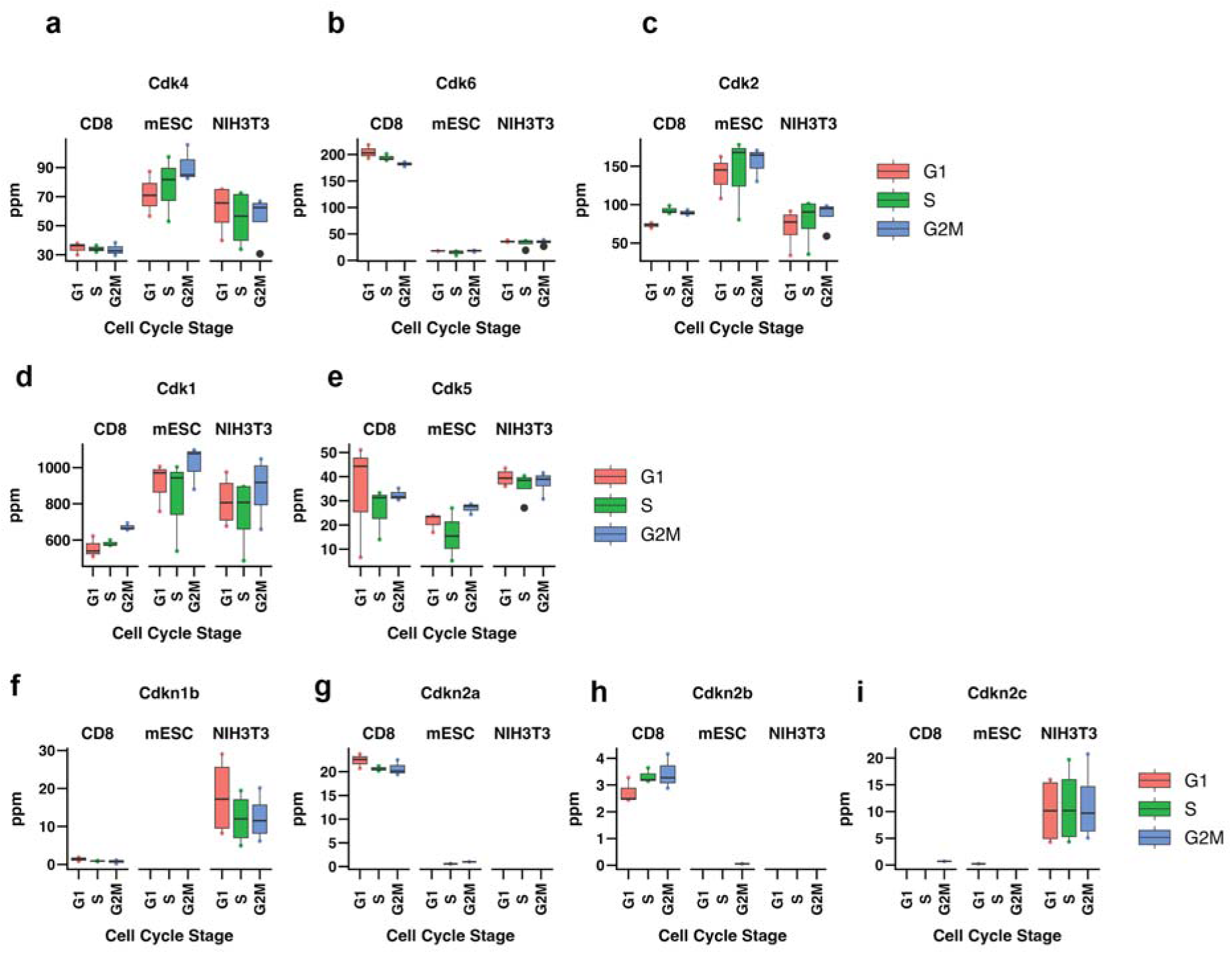
Comparing cell cycle regulation of Cdk and Cdk inhibitor proteins in fast and slow dividing cell types. **a-i** Protein intensities (ppm) for Cdk4 (a), Cdk6 (b), Cdk2 (c), Cdk1 (d), Cdk5 (e), Cdkn1b (p27) (f), Cdkn1a (p21) (g), Cdkn2b (p15) (h), and Cdkn2c (p18) (i).

While the majority of CDKs were not cell cycle regulated in abundance, CDKs showed cell type enhanced expression. For example, Cdk4 was higher expressed in mESC (Fig. 4a) whereas Cdk6 was higher expressed in CD8 T cells (Fig. 4b). These results suggest that the predominant complex between Cdk4/6 and D-type cyclins is Cdk6-Cyclin D3 in CD8 T cells and Cdk4-Cyclin D3 in mESC.

mESC also expressed higher levels of Cdk2 (Fig. 4c). Cdk1 levels were lower in CD8 T cells compared to the other two cell types (Fig. 4d). Interestingly, Cdk5 was higher expressed in CD8 T cells compared to mESC (Fig. 4e). Cdk5 was previously shown to be active in CD8 T cells and required for full TCR activation^44^. The cognate cyclin-like proteins for Cdk5 (p35/p39) were not detected in this dataset. However, recent results suggest that Cdk5 can form complexes with B-type cyclins^45^, which were expressed in CD8 T cells (Fig. 3c-d).

CDKs are inactivated by expression and binding to CDK inhibitor (CKI) proteins. We found cell type enhanced expression of CKIs. For example, p27 (Cdkn1b) was detected at low levels in all cell cycle phases in NIH3T3 cells and CD8 T cells (Fig. 4f). In contrast, p27 was not detectable in mESC. p16 (Cdkn2a) and p15 (Cdkn2b) are higher expressed in CD8 T cells (Fig. 4g-h), whereas p18 (Cdkn2c) was higher expressed in NIH3T3 (Fig. 4i). Comparing CKI levels to CDK and cyclin proteins, however, suggests that CKIs are likely to be present in relatively low, substoichiometric levels.

We conclude that the faster dividing CD8 T and mESC cell types are characterized by higher levels of canonical cell cycle promoting cyclins (A, B, D, E). However, except for Cyclin E1, the remaining cyclins fluctuated in abundance across the cell cycle. Interestingly, there were striking differences between mESC and CD8 T cells, including cell type-enhanced expression of CDKs, and higher levels of cell cycle regulated proteins in G1 phase in mESC (Fig. 2d).

### APC/C regulation in CD8 T cells

Numerous cell cycle regulated proteins are targeted for degradation in late mitosis and G1 phase by the APC/C E3 ligase^23,46,47^, including Cyclin A2 and Cyclin B1. It has been previously suggested that cell type-specific regulation of the APC/C E3 ligase underpins subdued periodicity in cyclin proteins in embryonic stem cells^3^. Consistent with this idea, in NIH3T3 and CD8 T cells, the abundances of common cell cycle regulated proteins are low in G1 phase, whereas mESC has higher levels (Fig. 2d). We next investigated if APC/C subunits and substrates showed cell-type specific expression.

The APC/C is a multi-subunit protein complex^48^ that requires binding to one of the two substrate adaptor proteins, Cdc20 or Cdh1 (*Fzr1*), for substrate ubiquitination. Out of the 14 core subunits, 10 were detected by mass spectrometry (Fig. 5a). 9 out of the 10 detected subunits were present at similar levels (2-fold or less) across cell type: Apc1 (*Anapc1*), Apc2 (*Anapc2*), Apc3 (*Cdc27*), Apc4 (*Anapc4*), Apc5 (*Anapc5*), Apc6 (*Cdc16*), Apc7 (*Anapc7*), Apc8 (*Cdc23*), and Apc16 (*Anapc16*). However, Apc10 (*Anapc10*) was significantly lower in all cell cycle phases measured in CD8 T cells compared to mESC and NIH3T3 cells (Fig. 5a, Fig. S1a), and Anapc16 and Cdc16 also showed lower expression (Fig. 5a). In contrast to core APC/C subunits, the levels of the co-activators showed significant differences between cell cycle phases and cell types. Cdc20 showed highest levels in G2&M phase compared to G1 and S phases (Fig. 5b), as previously shown^35,36,49^. Cdh1 (*Fzr1*) levels were low in CD8 T cells and NIH3T3 but markedly higher in mESCs (Fig. 5c).

**Figure 5:**
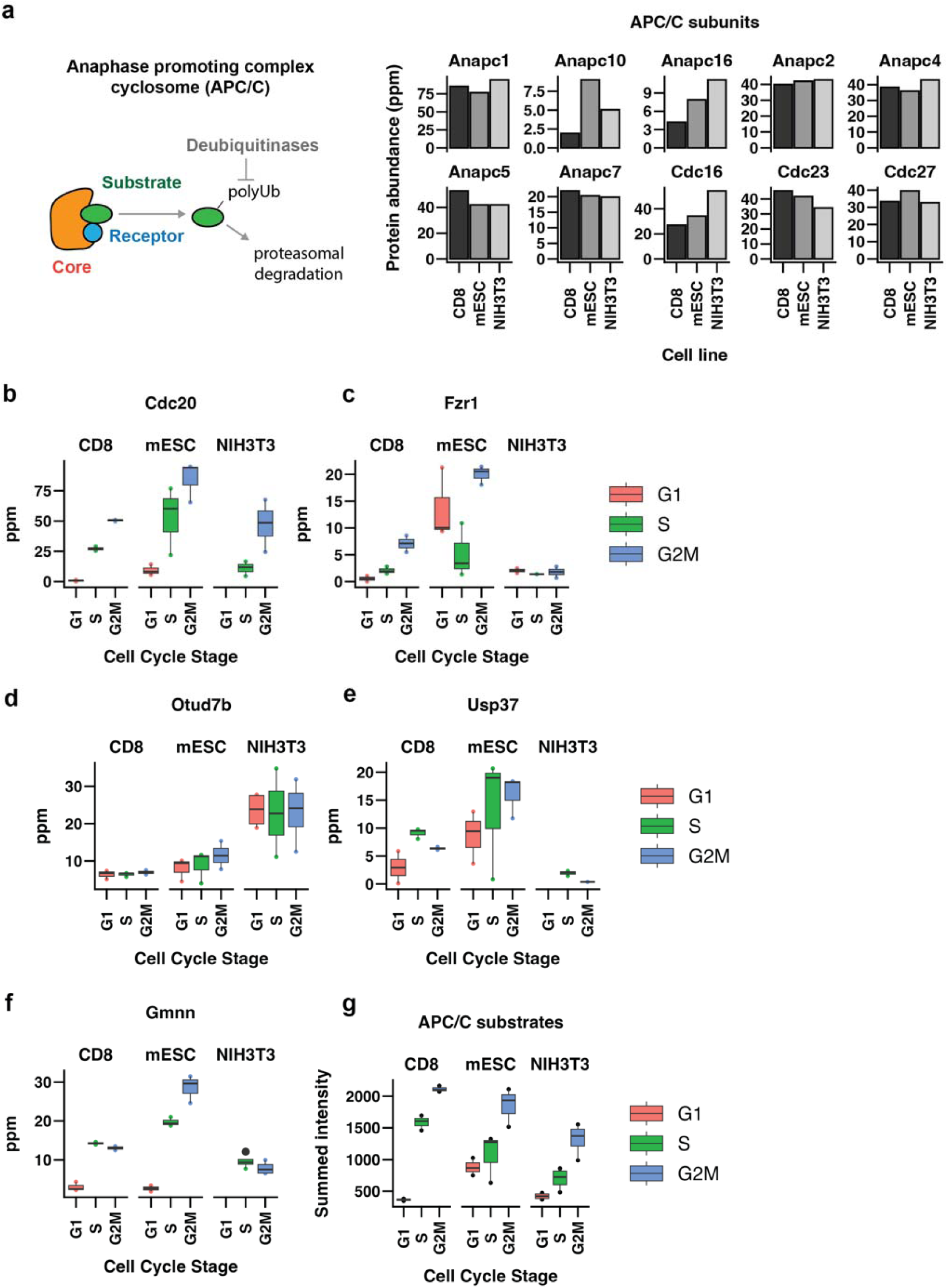
Comparing cell cycle regulation of the anaphase protein complex/cyclosome (APC/C) and its regulators. **a** Scheme illustrating APC/C molecular function (left) and protein intensities (ppm) measured for detected APC/C subunits. **b-f** Protein intensities (ppm) for Cdc20 (b), Fzr1 (c), Otud7b (d), Usp37 (e), and Gmnn (geminin) (f). **g** Summed protein intensities for APC substrates^53^ detected by proteomics (Supplementary Table 4).

APC/C substrate ubiquitination can be reversed by deubiquitinases (DUBs) that stabilize APC/C substrates and therefore promote cell cycle progression^50^. Two DUBs have been shown to antagonize APC/C ubiquitination, Otub7b/Cezanne^51^ and Usp37^52^, and were detected by MS (Fig. 5d-e). Otub7b was not cell cycle regulated in any of the three cell lines (Fig. 5d). However, NIH3T3 showed the highest levels of Otub7b protein. Usp37 abundance was cell cycle regulated in all three cell lines and in general, showed the highest levels in CD8 T cells and mESC (Fig. 5e). Interestingly, Usp37 levels were detected in G1 phase of the cell cycle in CD8 T cells and mESC (Fig. 5e). This would be consistent with a model whereby APC/C substrates were stabilized in early G1 to promote fast progression into S phase.

The previously proposed models suggested that APC/C activity is reduced in G1 phase, resulting in higher levels of APC/C substrates^1,3^. These models, however, are only partially consistent with our data. For example, median G1 expression of the 121 common cell cycle regulated proteins was higher in mESC. However, the APC/C substrates Cyclin A2 and Cyclins B1/B2 were not expressed, or present in similarly low levels in G1 phase across the three cell types examined. Indeed, the well characterized APC/C substrate Geminin (Gmnn) showed similar behaviour to Cyclins A2 and B1/B2 (Fig. 5f). Out of 33 validated APC/C substrates^53^, 26 were detected in this dataset (Supplementary Table 4). The abundances were summed as a readout of total levels of APC/C substrates. CD8 T cells and NIH3T3 showed similarly low APC/C substrate levels in G1 phase (Fig. 5g). In contrast, mESC showed a significantly higher level of APC/C substrates in G1. These results suggest that on average, APC/C substrates are maintained at higher levels in G1 phase of mESCs, with the best characterized APC/C substrates, namely Geminin and Cyclins, as striking exceptions. CD8 T cells do not share this behaviour despite their similarly fast cell cycles. The APC/C substrates with the highest abundance in mESC compared to NIH3T3/CD8 T cells are Tpx2, Ckap2, Nusap1, Kif11 (Eg5) and Kif22 (Fig. S1b). There is no consistent pattern in subcellular localization that could explain the differences observed in G1 protein abundance. For example, Tpx2, Nusap1 and Kif22 are localized to the nucleoplasm and Kif11 is localized to microtubules in interphase^54^. There are however functional associations between these proteins as Tpx2, Ckap2, Nusap1, Kif11 and Kif22 all bind to microtubules and/or regulate microtubule dynamics.

### Emi1 promotes G1/S progression in CD8 T cells

Emi1 is implicated in facilitating rapid mESC cell divisions by inhibiting the APC/C and stabilizing cell cycle promoting proteins, including Cyclin A2^3^ (Fig. 6a). While we did not observe increased Cyclin A2 levels in G1 phase in mESCs (Fig. 3a), the levels of other APC/C substrates were higher in G1 phase in mESC and not CD8 T cells or NIH3T3 cells (Fig. 5g). Our proteomics analysis showed that Emi1 was detected in G1 phase in CD8 T cells, but not in mESCs or NIH3T3 cells (Fig. 6b). Emi1 was also not detected in published proteomic analyses of human NB4 promyelocytes^35,37,49^, human TK6 lymphoblasts^36^ and human HeLa S3 epithelial cells^55^. While the abundance of APC/C substrates in G1 phase was highest in mESC (Fig. 5g), CD8 T cells also showed higher abundance of some APC/C substrates relative to NIH3T3 (Fig. S1b), consistent with a potential role of Emi1. We therefore next tested if Emi1 is functionally important in G1 progression and S-phase entry in CD8 T cells.

**Figure 6:**
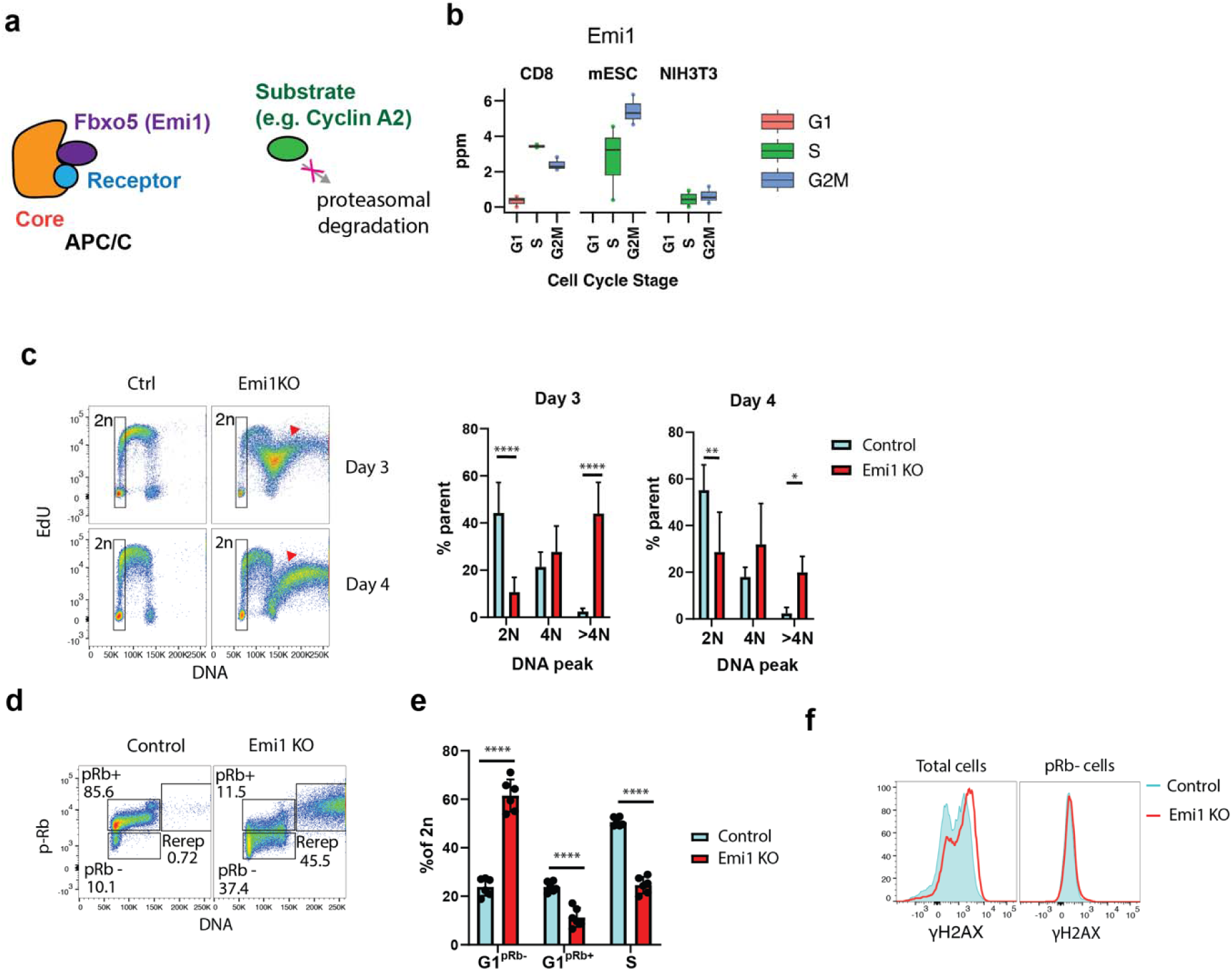
Emi1 promotes S-phase entry and safeguards against re-replication. **a** Scheme illustrating Emi1/Fbxo5 molecular function. **b** Protein intensities (ppm) for Emi1. **c** Activated CD8 T cells were transduced with the Cas9 complex with Emi1 targeted sgRNA or a blank control on day 2, cultured IL-2, and collected on Day 3 and Day 4 for analysis. Prior to harvest, cells were given a 30 minutes pulse of EdU. Cells were analyzed by flow cytometry. Frequency of cells containing >4N DNA. **d** Representative flow cytometry staining of phosphorylated Rb (residues S807/811) comparing control cells and Emi1KO. **e** Frequency of early G1/G0 (G1pRb-), late G1 (G1pRb+) and early S-phase cells at day 4 within the 2N DNA content gate as indicated in (b) **f** γH2AX expression of day 4 cells of the entire profile or of pRb-cells within the 2n gate. All flow data was gated to exclude the dead population and doublets. DNA staining was determined by Hoechst. Statistical significance was determined by unpaired Student’s t-test, ** p<0.01, *** p<0.001, **** p<0.0001, n = 6 mice from two independent cohorts.

The Emi1 locus was disrupted in activated CD8 T cells by CRISPR/Cas9 using guide RNAs (gRNAs) targeting exons 2 and 4, resulting in Emi1^CRISPR-KO^ CD8 T cells (hereafter called Emi1KO cells). Samples were collected for analysis 24 h and 48 h following targeted gene disruption. Emi1KO cells had indel mutations in the *Emi1* locus and reduced Emi1 protein levels (Fig. S2a-b). Consistent with a role of Emi1 in cell cycle progression, proliferation was significantly reduced in Emi1KO cells (Fig. S2c).

We next investigated the cell cycle phenotype in more detail. Emi1KO cells underwent DNA re-replication (Fig. 6c, Fig. S2d), as indicated by the presence of cells with >4N DNA content (Fig. S2d) that were also EdU+ (red arrowheads, Fig. 6c). Emi1KO cells were also larger in cell size compared to control cells, as inferred from forward scatter measurements (Fig. S2e). DNA re-replication and increased cell size are phenotypes observed following Emi1 depletion in transformed human cell lines^56,57^, thus validating the CRISPR approach in CD8 T cells.

To investigate the role of Emi1 specifically in G1/S, we excluded cells with >4N DNA and co-stained for pRb to identify pre- and post-restriction point G1 cells (G1^pRb-^ and G1^pRb+^). In cells that have not undergone DNA re-replication, Emi1KO increased the frequency of pRb^-^ from 10% in control cells to 37% in Emi1KO cells (Fig. 6d). Indeed, in cells with 2N DNA content (gate shown in Fig. 6c), there was an increase from 23% G1^pRb-^ in control cells to 60% G1^pRb-^ in Emi1KO cells, and corresponding decreases in G1^pRb+^ and early S-phase (EdU+) (Fig. 6e). To examine the possibility that increased frequency of G1^pRb-^ is due an abortive S-phase and return to G1 following DNA damage, we stained for the DNA damage marker gammaH2A.x. In cells ungated for cell cycle phase, we observed an increase in gammaH2A.x staining, consistent with DNA damage induced by Emi1KO in S-phase cells (Fig. 6f). However, G1^pRb-^ cells showed no difference in gammaH2A.x staining between control and Emi1KO cells. This suggests that the increased G1^pRb-^ and decreased S-phase frequencies observed in Emi1KO cells is likely due a function of Emi1 in promoting G1 progression and S-phase entry.

Together, these observations support the model that like in mESCs, Emi1 facilitates cell cycle progression in CD8 T cells.

### Cell cycle promoting factors are similarly expressed in cycling memory-like and effector CD8 T cells

CD8 T cells are multipotent^58^. Following activation, CD8 T cells can differentiate into various effector and memory phenotypes that have functions in pathogen elimination and long-term immunity, respectively. Differentiation can be promoted *in vitro* by culturing activated CD8 T cells with the mitogenic cytokines IL-2 for effector cells and IL-15 for “memory-like” cells (Fig. 7a). Compared to IL-2 treated cells, IL-15 treated cells have a higher frequency of G1^pRb-^ cells (Supp. Fig. 3a), suggesting that IL-15 treated memory-like cells have a slower cell cycle, consistent with previous reports^12^.

**Figure 7:**
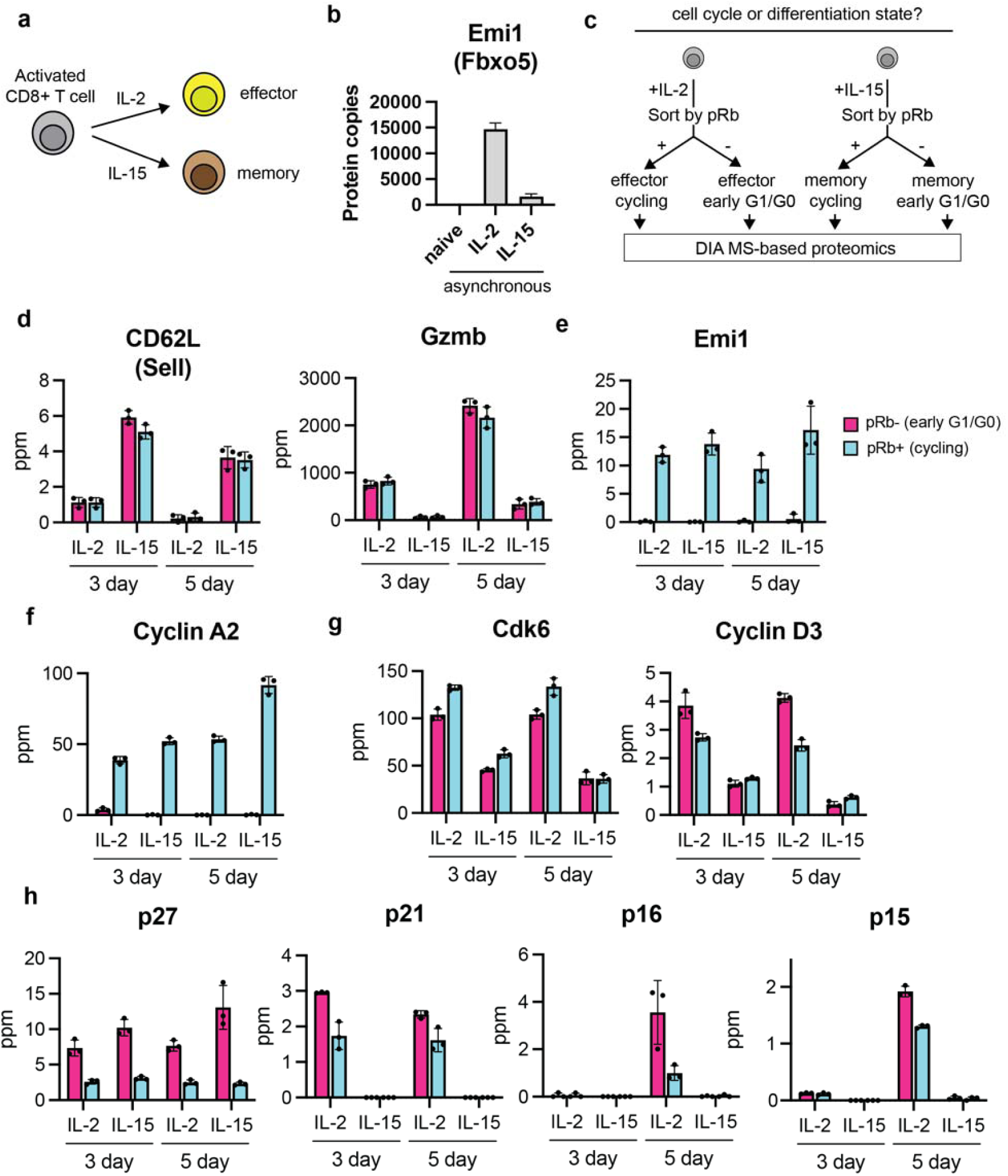
Emi1 is cell cycle regulated and promotes S-phase entry in both CD8 T effector and memory-like cells *in vitro*. **a** Schematic showing CD8 T cell phenotype specification in vitro by the IL-2 and IL-15 cytokines. **b** Emi1/Fbxo5 protein copies measured in activated CD8 T cells cultured in IL-2 versus IL-15 (cite ImmPRES). **c** Schematic illustrating approach to compare cell cycle regulated protein networks in effector and memory-like CD8 T cells. OT-1 CD8 T cells were activated with antigen (N4 peptide) for 2 days, and then cultured for an additional 2 days with indicated cytokine (IL-2 or IL-15) prior to harvest and analysis by data-independent acquisition (DIA) MS-based quantitative proteomics. **d** Protein abundance (ppm) of selected CD8 T cell phenotypic markers CD62L (Sell) and granzyme B (Gzmb). **e** Protein abundance (ppm) of Emi1/Fbxo5. **f** Protein abundance (ppm) of positive regulators of G1/S transition, Cdk6 and Cyclin D3. **g** Protein abundance (ppm) of negative regulators of cell cycle progression, including members of the CIP/KIP family (p21, p27) and INK4 family (p16, p15).

Interestingly, levels of Emi1 were significantly lower in asynchronous IL-15 treated cells compared to asynchronous IL-2 cells (Fig. 7b)^59^. Because Emi1 facilitated G1 progression (Fig. 6), it is possible that lower Emi1 levels resulted in slower cell division cycles in IL-15 treated cells. Alternatively, it is equally possible that because Emi1 levels are cell cycle regulated (Fig. 6b), the difference in Emi1 levels could be explained by differences in cell cycle phase frequency following prolonged culture (Fig. S3a). To distinguish between these possibilities, IL-2 and IL-15 treated cells were FACS separated into pRb^-^ and pRb^+^ populations, which correspond to early G1/G0 and cycling cells, respectively, and total protein levels were measured proteome-wide (Fig. 7c). CD62L, also known as L-selectin and encoded by the *Sell* gene is a marker of memory phenotype and was highest in IL-15 treated cells (Fig. 7d). Granzyme B (Gzmb) is a marker of effector phenotype and was highest in IL-2 treated cells (Fig. 7d). These results are therefore consistent with IL-2 and IL-15 driving effector and memory-like differentiation, respectively. Emi1 levels, however, were consistently low in pRb^-^ cells and high in pRb^+^ cells, irrespective of cytokine treatment (Fig. 7e). This suggests the difference in Emi1 levels observed in asynchronous IL-15 versus IL-2 treated cells (Fig. 7b) likely reflected a change in the cell cycle phase distribution promoted by cytokine treatment.

These observations raised the question of whether Emi1 functions similarly in IL-15 treated cells to promote G1 progression. IL-15 treated Emi1KO cells induced re-replication (Fig. S3b-d), an increase in cell size (Fig. S3e), and increased G1^pRb-^ cell frequency (Fig. S3f), resembling the phenotype of Emi1KO IL-2 treated cells shown in Fig. 6. This similarity suggests that Emi1 acts downstream of the critical factors that control cell cycle entry following differential cytokine treatment and instead promotes rapid S-phase entry post-commitment.

Interestingly, like Emi1, A- and B-type cyclins fluctuated by similar or even higher fold-changes in IL-15 versus IL-2 treated cells (Fig. 7f, Fig. S3g). These factors include Cyclin A2, Cyclin B1 and Cyclin B2 and other proteins that promote cell cycle progression. These data suggest that once cells become committed to cell cycle progression, i.e., becoming pRb^+^, IL-15 treated cells spend a similar time to IL-2 treated cells completing a cell division cycle. Cdk6 and Cyclin D3 were notable exceptions to the general trend exemplified by Cyclin A2. Both Cdk6 and Cyclin D3 were higher in IL-2 compared to IL-15 treated cells (Fig. 7g).

Examination of CDK inhibitor proteins of the CIP/KIP (CDK interacting protein/kinase inhibitory protein) and INK4 (inhibitor of CDK4) family showed differential expression patterns. The CIP/KIP protein p27 was higher in pRb^-^ compared to pRb^+^ cells irrespective of cytokine treatment. This lack of cytokine-dependence for p27 resembles other core cell cycle proteins, like Emi1 and Cyclin A2. However, expression of p21, p16, and p15 was specific to IL-2 treated cells. p16 and p15 are associated with terminal cell cycle exit and cellular senescence^60^.

Interestingly, p16 and p15 increased in expression with time in culture with IL-2 (Fig. 6h), suggesting that CD8 T cells may acquire senescent-like phenotypes following prolonged culture in IL-2.

### Emi1KO modulates the immune phenotype of activated CD8 T cells

The results above suggest that differentiation state imposed by cytokine treatment affects cell cycle distribution (Fig. S3a) and the abundance of G1/S regulators (D-type cyclins and INK4 family proteins). We next aimed to investigate if a causal relationship existed in the opposite direction, and specifically if disrupting cell cycle progression by Emi1KO affects CD8 T cell immune phenotype.

To begin investigating this, we examined markers of immune phenotype. CD62L is important for the tissue trafficking of lymphocytes to secondary lymphoid organs and is therefore frequently used as a marker of CD8 T cell central memory phenotype in combination with other markers^61^. Consistent with this, increased CD62L protein was detected in IL-15-treated cells compared to IL-2 treated cells (Fig. 7d). CD25 (encoded by the *Il2ra* gene) is the alpha subunit of the high affinity IL-2 receptor and is upregulated following TCR stimulation. CD25 is frequently used as a marker of activated CD8 T cells. We tested whether perturbing cell cycle progression by disrupting Emi1 gene function affects the frequency and magnitude of CD62L and CD25 expression.

Interestingly, Emi1KO resulted in a significant increase in the frequency of CD62L^+^ cells in both IL-2 and IL-15 treated cells (Fig. S4a) and magnitude of CD62L expression (Fig. S4b), suggesting that Emi1 plays a role in regulating CD62L. Emi1KO produces a slight but significant increase in CD25 expression in IL-2 treated cells (Fig. S4c). An increase was also observed in IL-15 treated cells but was not significant (Fig. S4c). CD62L levels (Fig. 7d) and CD25 levels (Fig. S4d) are not cell cycle regulated (Supplementary Table 2), suggesting that the effects observed on CD62L and CD25 are a consequence of Emi1 loss of function, rather than cell cycle per se. We conclude that Emi1KO significantly affects CD8 T cell phenotype by modulating CD62L frequency and expression.

## Discussion

The structure and control of the cell division cycle are developmentally plastic to facilitate the differentiation of embryonic stem cells into somatic cell types that vary in propensity for proliferation. Unlike many other cells in an adult mammal^62^, T cells retain high proliferation potential and undergo several rounds of rapid proliferation followed by dormancy during thymic development and adaptive immunity. The mechanisms T cells employ to promote rapid proliferation remain poorly understood. Here, we have focused on the rapid cell cycles of mature, stimulated CD8 T cells.

We have provided a quantitative comparison of cell cycle regulated protein abundance comparing CD8 T cells, mESCs and NIH3T3 fibroblasts that are conspecific cell lines that vary in cell cycle speed. To do this, we have used PRIMMUS^35,36^, an approach that avoids arrest-based synchronization that has the potential to promote differentiation of multipotent cells. Furthermore, PRIMMUS avoids stress-induced artefacts that are not related to cell cycle per se. We identified 121 proteins that are common cell cycle regulated proteins. Interestingly, many of these proteins in general showed higher abundance in the faster dividing cells (CD8 T cells, mESCs) compared to slower dividing NIH3T3 cells. Of the cell cycle cyclins and CDKs examined, only Cyclin E1 showed constitutive expression across the cell cycle in mESCs and CD8 T cells. This is consistent with earlier work showing that Cyclin E-Cdk2 activity is constant in ESC and becomes periodic following loss of pluripotency. Our results suggest that CD8 T cells also utilize Cyclin E1 to drive rapid immune cell clonal expansion.

mESCs had a higher abundance of APC/C substrates in G1 phase of the cell cycle, consistent with previous reports that APC/C activity during G1 is attenuated. Curiously, APC/C substrates differ in their abundance in G1. We found that well characterized APC/C substrates including A- and B-type Cyclins and Geminin are at very low levels or undetectable in G1 phase in the three cell lines examined. In contrast, several other APC/C substrates are present at relatively high levels in G1 phase specifically in rapidly dividing cells. These proteins include Tpx2, Ckap2, Nusap1, Kif11, and Kif22, which are all microtubule binding proteins. In contrast, APC/C substrates are generally present in very low levels in CD8 T cells. Interestingly, specific APC/C substrates showed higher abundance in G1 phase of CD8 T cells (e.g. Kif22, Fig. S1b). It will be important in future to investigate whether APC/C substrates are subject to selective stabilization in a cell type dependent manner.

Supporting a role of APC/C in CD8 T cell cycle control, the levels of its inhibitor, Emi1, were high in CD8 T cells and disruption of Emi1 function by CRISPR/Cas9 showed defects in G1 progression and S-phase entry in CD8 T cells. It is still possible that Emi1KO results in low level DNA damage in cells with 2N DNA content that cannot be detected using gammaH2A.x immunofluorescence, and therefore further work is required to better understand the role of Emi1 in rapid cell cycles. Nevertheless, these results suggest that like mESCs, Emi1 is important to promote cell cycle progression in CD8 T cells. Additionally, it will be important to determine which proteins are affected by Emi1 deletion in CD8 T cells and mESCs to understand if increased levels of these proteins are due to Emi1-medated inhibition of APC/C.

Constitutive expression of specific cell cycle regulators has been proposed as a mechanism to promote rapid proliferation in early embryonic cell cycles^1^. In contrast to previous reports that Cyclin A2 is either constitutively expressed, or has dampened periodicity in mESCs, our data show that Cyclin A2 levels are highly periodic in the three cell lines examined. Our results suggest that differences in Cyclin A2 regulation are unlikely to explain variation in cell cycle duration. Our results instead suggest that rapid cell divisions may be promoted by constitutive expression of Cyclin E1, reduced levels of CDK inhibitor proteins, and quantitatively higher levels of A-, B- and D-type Cyclin proteins. Faster dividing cells also express higher levels of the Usp37 deubiquitinase and Emi1, both proteins that antagonize or directly inhibit APC/C-mediated degradation of cell cycle promoting proteins. These data point to multiple mechanisms to support rapid cell cycles and further work is needed to understand the relative importance of these mechanisms.

Our data on CD8 T cells support an emerging paradigm whereby cell cycle control pathways can be targeted to modulate the immune system. In this study, we showed that genetic disruption of the cell cycle regulator, Emi1, increases the frequency of G1 and CD62L+ cells, with CD62L a marker associated with CD8 T central memory cells. This is consistent with an idea that delaying G1/S promotes memory formation, as suggested previously^29^. Further investigation into the mechanism is warranted to separate the function of Emi1 in G1/S from its function in safeguarding against DNA re-replication. It will also be important to understand how other chemotherapeutics that target G1/S progression, for example recently described CDK2-specific^63–65^ and CDK2/4/6 inhibitors^64,66^, affect immune function.

Previous studies have suggested that memory-like CD8 T cells cycle more slowly. Consistent with this idea, cell cycle regulators are lower in abundance in asynchronous IL-15-treated cells compared to IL-2 treated cells, as illustrated for Emi1 (Fig. 7b). In this study, we fractionated cells into early G1/G0 cells (pRb-) and cycling (pRb+) cells and showed that cell cycle regulators in IL-15 treated cells are present in similar, if not higher abundance, to IL-2 treated CD8 T cells. Our results support an alternative, digital model of cell cycle entry in CD8 T cells. IL-15 treated cells have a decreased cell cycle entry frequency, leading to the differences observed in this study and previously. It will be important to investigate the cycling time of cell cycle committed IL-15 treated CD8 T cells, which we predict based on the abundances of cell cycle regulators, will be similar, if not faster than IL-2 treated cells. Curiously, Cyclin D3 and Cdk6 are lower in IL-15 treated cells, relatively independent of cycling status (Fig. 6f), and regulate the classic cell cycle ‘commitment’ point (or restriction point). This pathway would be a candidate for inhibiting cell cycle entry specifically in memory-like CD8 T cells.

## Acknowledgements

We thank members of the Ly, Zamoyska, and Doreen Cantrell groups for advice and feedback. This work is supported by a Wellcome Trust and Royal Society Sir Henry Dale Fellowship to T.L. (206211/Z/17/Z); an EASTBIO PhD studentship to D.A.L.; a Wellcome Multi-User Equipment Grant to T.L. (218305/Z/19/Z); a BBSRC grant to T.L. (BB/X007057/1); and a Wellcome grant to R.Z. (WT205014/Z/16/Z).

## Methods

### Ethics Statement

This study was approved by the Ethical Review Body at the School of Biological Sciences, University of Edinburgh. All animal experiments were approved by the University of Edinburgh Bioresearch and Veterinary Services Ethical Review body and the United Kingdom Home office under project licence P38881828 to R.Z.

### Mice

Mice expressing the OT-1 TCR transgene (C57BL/6-Tg(TcraTcrb)1100Mjb/J) backcrossed to the Rag-1KO (B6.129S7-*Rag1^tm1Mom^*/J) background (C57BL/6 Rag-1KO OT-1 mice) were used as lymph node donors for CD8 T cells. All mice were bred and housed in individually ventilated cages under specific pathogen-free conditions at the University of Edinburgh Bioresearch and Veterinary Services (BVS) facilities. OT-1 animals express a transgenic TCR specific for the ovalbumin peptide OVA257-264 (SIINFEKL), also known as N4 peptide.

Mice were between 5 and 12 weeks of age and sexes were randomised for tissue donors. Mice were maintained under a 12 h light/12 h dark cycle with ad libitum access to food and water at a temperature of 19–24 °C and humidity of 45–65%.

### Cell lines

hTERT-RPE1 cells were purchased from ATCC and cultured in DMEM/F-12 + 10% Fetal Calf Serum (FCS). NIH3T3 cells were purchased from ATCC and cultured in DMEM + 10% FCS. Mouse ES cells (E14tg2A) were maintained in DMEM (ThermoFisher Scientific, 11960044) supplemented with 10% FCS (FCS-SA/500, LabTech), 5% KnockOut Serum Replacement (ThermoFisher Scientific, 10828028), 2 mM L-Glutamine (ThermoFisher Scientific, 25030081), 100 U/ml Penicillin–Streptomycin (ThermoFisher Scientific, 15140122), 1 mM Sodium Pyruvate (ThermoFisher Scientific, 11360070), a mixture of seven non-essential amino acids (ThermoFisher Scientific, 11140050), 0.05 mM β-mercaptoethanol (Sigma-Aldrich, M6250) and 0.1 μg/ml Leukaemia Inhibitory Factor (MRC PPU Reagents and Services, DU1715). Cells were grown at 37°C in a humidified atmosphere of 5% CO_2_. For passaging, cells were released from dishes using 0.05% Trypsin–EDTA (ThermoFisher Scientific, 25300054).

### CD8 T cell culture and stimulation

OT-1 T cells were obtained by passing tissue through 70 μM mesh filters and were cultured in IMDM medium (Sigma-Aldrich I3390) supplemented with 10% FCS, L-glutamine, 100U/ml penicillin, 100U/ml streptomycin and 50 μM 2β-ME. Activation was induced by incubation with 6 nM N4 antigen peptide SIINFEKL (HY-P1771A-5mg; Cambridge bioscience) and 1:500 αCD28 (Cat:553294 Clone:37.51; BD Pharmigen) for up to 48 hours. Activated T cells were then washed in media and returned to culture with 20 µg/ml human IL-2 (200-02; Peprotech), or 10 µg/ml IL-15 (200-15; Peprotech) with daily supplements of equivalent volume cytokine added in 24 hour intervals for the duration of the culture.

### Flow cytometry

Cells were removed from culture and stained with Live/Dead fixable near-IR Dead cells stain kit (L34975;Thermofisher) as per manufacturer’s instructions.

Surface staining was done in FACS buffer (0.1% BSA (Fisher bioreagents; BP9702-100), 0.01% Sodium Azide (S8032; Sigma-Aldrich), PBS (D1408-500ml; Sigma-Aldrich) with antibodies at the appropriate concentration for 20 minutes at 4°c.

Cells were washed in PBS and resuspended and fixed in 2% Formaldehyde (28908; Thermofisher) for 30 minutes at room temperature in constant rotation.

Cells requiring intracellular staining were resuspended in <10 µl FACS buffer. Once resuspended, they were permeablised by 500µl ice cold 90% MeOH (M/4000/PC17; Fisher bioreagents) was added dropwise with gentle vortexing to the cells and stored at −20°C.

Prior to the intracellular stain, the cells were washed twice with FACS buffer and then incubated with antibodies at the appropriate concentration for 1 hour at 4°C. The cells were then washed and acquired in FACs buffer or FACs buffer with 1 µg/ml Hoechst 33342 Data were acquired using either a MACSQuant Analyzer (Miltenyi), or a Fortessa (BD Biosciences).

Formaldehyde fixed, methanol permeabilized CD8 T cells were stained with anti-phosphoRb (S807/S811) and Hoechst and separated into G1^2N,pRb-^, G1^2N,pRb+^, S^2N-4N,pRb+^, and G2&M^4N,pRb+^ phase fractions using a FACSAria IIu sorter (BD Biosciences). Pilot pRb staining performed on mESCs and NIH3T3 showed the absence of a pRb negative population. Therefore, for sorting, no pRb staining was performed. Formaldehyde fixed, methanol permeabilized mESCs and NIH3T3 were stained with EdU and Hoechst and separated into G1^2N,EdU-^, S^2N-4N,EdU+^ and G2&M^4N,EdU-^ phase fractions using a MA900 sorter (Sony Biosciences).

### Antibodies

| Specificity | Conjugate | Clone | Company | Cat. number |
| --- | --- | --- | --- | --- |
| m CD8a | PerCP | 53-6.7 | BioLegend | 100732 |
| m CD8a | AF647 | 53-6.7 | BioLegend | 100724 |
| m CD8b | PE-Cy7 | H35-17.2 | Invitrogen | 25-0083-82 |
| m CD25 | PE | PC61 | BioLegend | 102008 |
| m CD25 | PE-Cy7 | PC61 | BioLegend | 102016 |
| m CD25 | APC | PC61 | BioLegend | 102012 |
| m CD44 | ef450 | IM7 | eBiosciences | 48-0441-82 |
| m CD44 | APC-eF780 | IM7 | Invitrogen | 47-0441-82 |
| m CD69 | AF488 | H1.2F3 | BioLegend | 104516 |
| m CD69 | APC | H1.2F3 | BioLegend | 104514 |
| p-H3(Ser10) | PE | D7N8E | Cell Signaling | 29237S |
| p-Rb(Ser807/811) | AF488 | D20B12 | Cell Signaling | 4277S |
| Emi1 | AF546 | B-3 | Santa Cruz | Sc365212<br>AF546 |
| Eg5 | AF647 | EPR23276-52 | Abcam | ab283273 |
| γH2AX | PE | 20E3 | Cell Signaling | 5763S |
| Emi1 | (Rabbit Ab) | EPR15320 | Abcam | AB184950 |
| Zap70 | (Mouse Ab) | 29/Zap70 Kinase | BD Biosciences | 610239 |
| Mouse IgG | IRDye 800CW | Polyclonal Goat | Licor | 926-32210 |
| Rabbit IgG | AF680 | Polyclonal Donkey | Life Technologies | A10043 |

### EdU click reaction

Cells were stained for EdU with use of the Click-IT EdU imaging kit (C10337;Invitrogen) as per manufacturer’s instructions with some modifications. Cells in cultured were pulsed with 1 µM EdU for 30 minutes prior to removal of culture and processed for flow cytometry as above. Prior to intracellular staining, the cells were incubated with the click reaction mix, swapping out the azide-488 that came with the kit for azide-AF594 (CLK-1295-1; Jena Bioscience).

#### CellTraceViolet Proliferation assay

Naïve cells were washed with PBS and incubated at 37C for 20 minutes with 5 µM Cell Trace Violet (C34571A; Invitrogen) at a volume of 1ml per 1e7 cells. Following incubation, the cells were quenched with 2x volume of media and incubated at 37C for 5 minutes.

#### Immunoblot analysis

Cell pellets were lysed using 2% SDS + protease inhibitor cocktail (P8340ML; Sigma-Aldrich) incubating at 65C for 5 minutes, 1 µl Benzonase (70664-3; Sigma-Aldrich) was added and incubated for a further 15 minutes. The lysates were then sonicated with a Ultrasonic Processor (UP200ST; Hielscher) at full power for two cycles of 20s on 30s off. Protein content for lysates were then assessed using Micro BCA Protein Assay Kit (23235; Thermofisher) as per manufacturer’s instruction.

Lysates were incubated with NuPage LDS Sample Buffer (NP0007; Invitrogen) and incubated at 95°C for 5 minutes and loaded into NuPage 4%-12% Bis-Tris gradient MOPS gel (NP0323BOX; Invitrogen) in MOPS SDS Running Buffer (NP0001; Invitrogen) and run at 70 volts for 15 minutes and then 140 volts for 75 minutes. The gel was when blotted on to an Immobilon-FL PVDF membrane (IPFL00010; Millipore Sigma) submerged in transfer buffer (24mM Tris Base (BP152-1; Fisher Bioreagents), 192mM Glycine (G8898-1KG; Sigma-Aldrich), 20% Ethanol (20821.330; VWR)) at 120 volts for 105 minutes, and then left to block in Intercept Blocking buffer (927-70001; Licor) for 1 hour or overnight.

The membrane was stained in primary antibody diluted in blocking buffer + 0.1% Tween 20 (H5151; Promega) for 4 hours or overnight, and the secondary antibody diluted in blocking buffer 0.1% Tween 20 and 0.01% SDS. The membrane was washed following both stains in PBS + 0.1% Tween.

The membrane was imaged using a Licor Odyssey CLx (Licor) and analysed using ImageJ.

### Proteomics sample preparation

Sorted cells undergoing mass spec analysis were washed and resuspended in triethylammonium bicarbonate (TEAB) digestion buffer (100 nM TEAB, 50 µM CaCL_2_, 50 µM MgCL_2_), incubated with 1 µl Benzonase at 37C for 15-30 minutes, and then digested using the in-cell digestion protocol^36^ using Pierce Trypsin Protease (90057; Thermo Fisher) overnight at a ratio of 1 µg trypsin per 20 µg of protein. Peptides were acidified in either 1% trifluoroacetic acid (TFA). Fragments of C18 silica membrane were placed in 200 µl tips, wetted with 100% acetonitrile (MeCN) (A955-1; Fisher Chemicals) and primed with 1% TFA/formic acid (10596814; Thermo Fisher). For samples in excess of 100 µg, higher capacity Sep-Pak Vac tC18 cartridges (WAT036790; Waters) were used instead. The sample was loaded onto pre-conditioned columns, washed with 0.1% TFA, and then eluted into fresh tubes with 70% MeCN diluted in either 0.1% TFA, or 0.1% formic acid.

### Mass spectrometry

For data-independent analysis (DIA): Peptides (equivalent of 1 µg) were injected onto a C18 reverse-phase chromatography system (UltiMate 3000 RSLC nano, Thermo Scientific) and electrosprayed into an Orbitrap Exploris 480 Mass Spectrometer (Thermo Fisher). For liquid chromatography the following buffers were used: buffer A (0.1% formic acid in Milli-Q water (v/v)) and buffer B (80% acetonitrile and 0.1% formic acid in Milli-Q water (v/v). Samples were loaded at 10 μL/min onto a trap column (100 μm × 2 cm, PepMap nanoViper C18 column, 5 μm, 100 Å, Thermo Scientific) equilibrated in 0.1% trifluoroacetic acid (TFA). The trap column was washed for 3 min at the same flow rate with 0.1% TFA then switched in-line with a Thermo Scientific, resolving C18 column (75 μm × 50 cm, PepMap RSLC C18 column, 2 μm, 100 Å).

Peptides were eluted from the column at a constant flow rate of 300 nl/min with a linear gradient from 3% buffer B to 6% buffer B in 5 min, then from 6% buffer B to 35% buffer B in 115 min, and finally from 35% buffer B to 80% buffer B within 7 min. The column was then washed with 80% buffer B for 4 min. Two blanks were run between each sample to reduce carry-over. The column was kept at a constant temperature of 50C.

The data was acquired using an easy spray source operated in positive mode with spray voltage at 2.40 kV, and the ion transfer tube temperature at 250°C. The MS was operated in DIA mode. A scan cycle comprised a full MS scan (m/z range from 350-1650), with RF lens at 40%, AGC target set to custom, normalised AGC target at 300%, maximum injection time mode set to custom, maximum injection time at 20 ms, microscan set to 1 and source fragmentation disabled. The MS survey scan was followed by MS/MS DIA scan events using the following parameters: multiplex ions set to false, collision energy mode set to stepped, collision energy type set to normalized, HCD collision energies set to 25.5, 27 and 30%, orbitrap resolution 30000, first mass 200, RF lens 40%, AGC target set to custom, normalized AGC target 3000%, microscan set to 1 and maximum injection time 55 ms. Data for both MS scan and MS/MS DIA scan events were acquired in profile mode.

### Proteomic data analysis

RAW data files from DIA experiments were processed using Spectronaut (version 16.2.220903.53000, Biognosys, AG). The directDIA workflow, using the default settings (BGS Factory Settings) with the following modifications was used: decoy generation set to mutated; Protein LFQ Method was set to QUANT 2.0 (SN Standard) and Precursor Filtering set to Identified (Qvalue); Precursor Qvalue Cutoff and Protein Qvalue Cutoff (Experimental) set to 0.01; Precursor PEP Cutoff set to 0.1 and Protein Qvalue Cutoff (Run) set to 0.05. For the Pulsar search the settings were: maximum of 2 missed trypsin cleavages; PSM, Protein and Peptide FDR levels set to 0.01; scanning range set to 300-1800 m/z and Relative Intensity (Minimum) set to 5%; cysteine carbamidomethylation set as fixed modification and acetyl (N-term), deamidation (asparagine, glutamine), dioxidation (methionine, tryptophan), glutamine to pyro-Glu and oxidation of methionine set as variable modifications. The database used was M.musculus downloaded from uniprot.org on 2022-10-25 (55,311 entries).

The output from Spectronaut was then processed in R (4.0.3). Protein abundances were normalized for technical differences in peptide loading by dividing each protein intensity by the sum of all protein intensities and multiplying the result by 1e6 (parts per million, ppm). A ppm is thus equal to one part per million intensity units detected in the sample. G1 intensities in the CD8 T cell dataset were corrected for S-phase contamination by calculating the %EdU+ cells in the 2N DNA content gate (e.g. Fig 1c). This fraction was then multiplied by the S-phase intensity to estimate the fractional intensity originating from S-phase contamination and subtracted from the intensity measured in the G1 fraction. GO term analysis was done by exporting a list of proteins from the dataset and retrieving the Gene Ontology (GO) terms from Uniprot.org. Further analysis of GO terms was done utilizing the DAVID Bioinformatics resources (https://david.ncifcrf.gov/)^67^. To identify cell cycle regulated proteins in all three cell lines, first missing values were imputed using Gaussian-based distributions for noise^68^. Fold changes were calculated and ANOVAs performed to identify significantly changing proteins in each cell line using the following filtering criteria: p < 0.05 and fold change > 2. These proteins were then clustered using hierarchal clustering using the “complete” agglomeration algorithm implemented in R. The cluster that showed cell cycle regulated protein abundance in all three cell lines was used for further analysis.

### CRISPR

Target PAM sites for CRISPR were identified using ChopChop (https://chopchop.cbu.uib.no/) with default settings^69^. Candidates were selected by for their high GC content, low number of mis-match pairs, and high efficiency. Candidate gRNA targeting exon 2 and exon 4 were synthesized (Integrated DNA Technologies).+

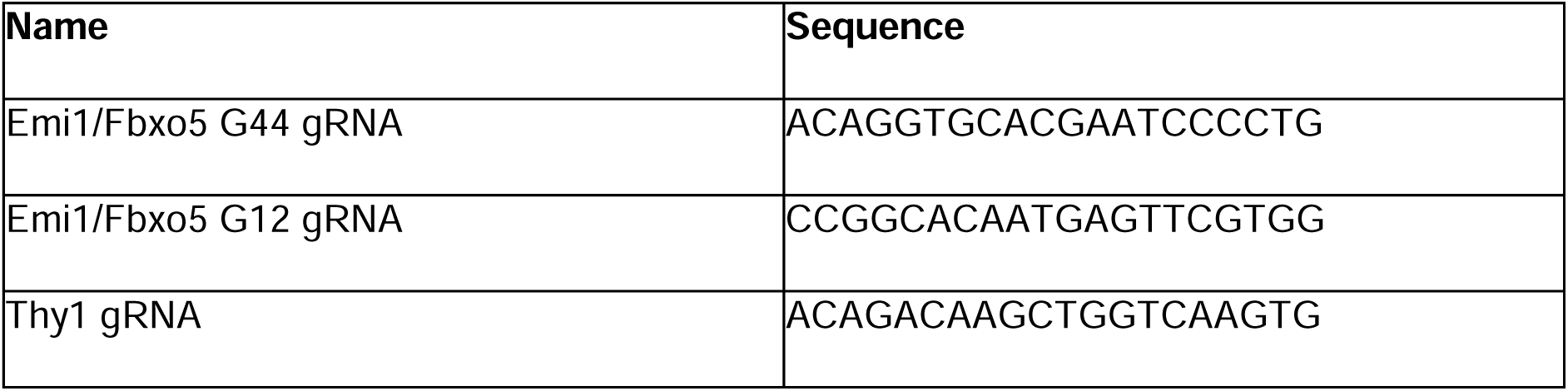

gRNA-Cas9 transfection was performed using Neon Transfection 100 µl kit (MPK10096; Invitrogen). gRNA was incubated with tracrRNA (B00817; IDT) in Nuclease free duplex buffer (1072570; IDT) for 5 minutes at 98C. Samples were then allowed to cool down before incubating with the TrueCast Cas9 protein V2 (A36499; Thermo Fiasher) and Resuspension Buffer T for 20 minutes at 37C. T cells were pelleted at 300G for 5 minutes and resuspended in Resuspension Buffer T at a concentration of 1.2e6 cells/100ul. 1.2e6 cells were mixed with the assembled Cas9 complex and electroporation was carried out utilizing the Neon 100 µl Tips within a solution of Electrolytic Buffer E2 provided by the kit. Electroporation was induced using the Neon Transfection System (Invitrogen) with a programme of 3 pulses of 10 ms 1600V for Activated T cells and 4 pulses of 5 ms 2500V for naive T cells.

### TIDE analysis

Primers were designed to assess CRISPR KO efficiency via TIDE^70^. The criteria for their design were: 1) the primer target sequence must be between 250-450bp from the Cas9 cut site, and 2) that the PCR product size must be ≥ 500bp long. These criteria were implemented in the Primer-BLAST webtool (https://www.ncbi.nlm.nih.gov/tools/primer-blast/), and primers chosen based on fewest number of off-target matches.

PCR mixes were created using the Phusion High-Fidelity DNA Polymerase Reagents (M0530L; New England Biolabs). Samples were spun down and removed of liquid before being lysed with PCR Lysis buffer (0.1% SDS, 0.05% Tween, 400ug/ml Proteinase K (25530-015; Life Technologies). PCR was then conducted with the following settings on a T100 Thermo Cycler (Biorad). 98C for 1 minutes, 98C for 15 seconds, optimal temperature (between 62C and 72C) for 15 seconds, 72C for 45 seconds, cycle 34 times, then 72C for 10 minutes, and hold at 12C. A fraction of the PCR product was run through a 1.5% Agarose gel made from UltraPure Agarose (16500-500; Invitrogen) in TAE buffer. Electrophoresis was performed at 120V for 20 minutes and imaged on a Licor D-Digit. The remaining product was then purified using the Monarch PCR & DNA Cleanup Kit (T1030L;New England Biolabs). The purified product was then quantified with a Qubit 3.0 (Invitrogen) and sent to the MRC Protein Phosphorylation and Ubiquitylation Unit in Dundee (https://dnaseq.co.uk/) for sequencing. The returned sequences were then analysed using the TIDE analysis webtool (https://tide.nki.nl/)^70^.

---

**Supplementary Figure 1:**
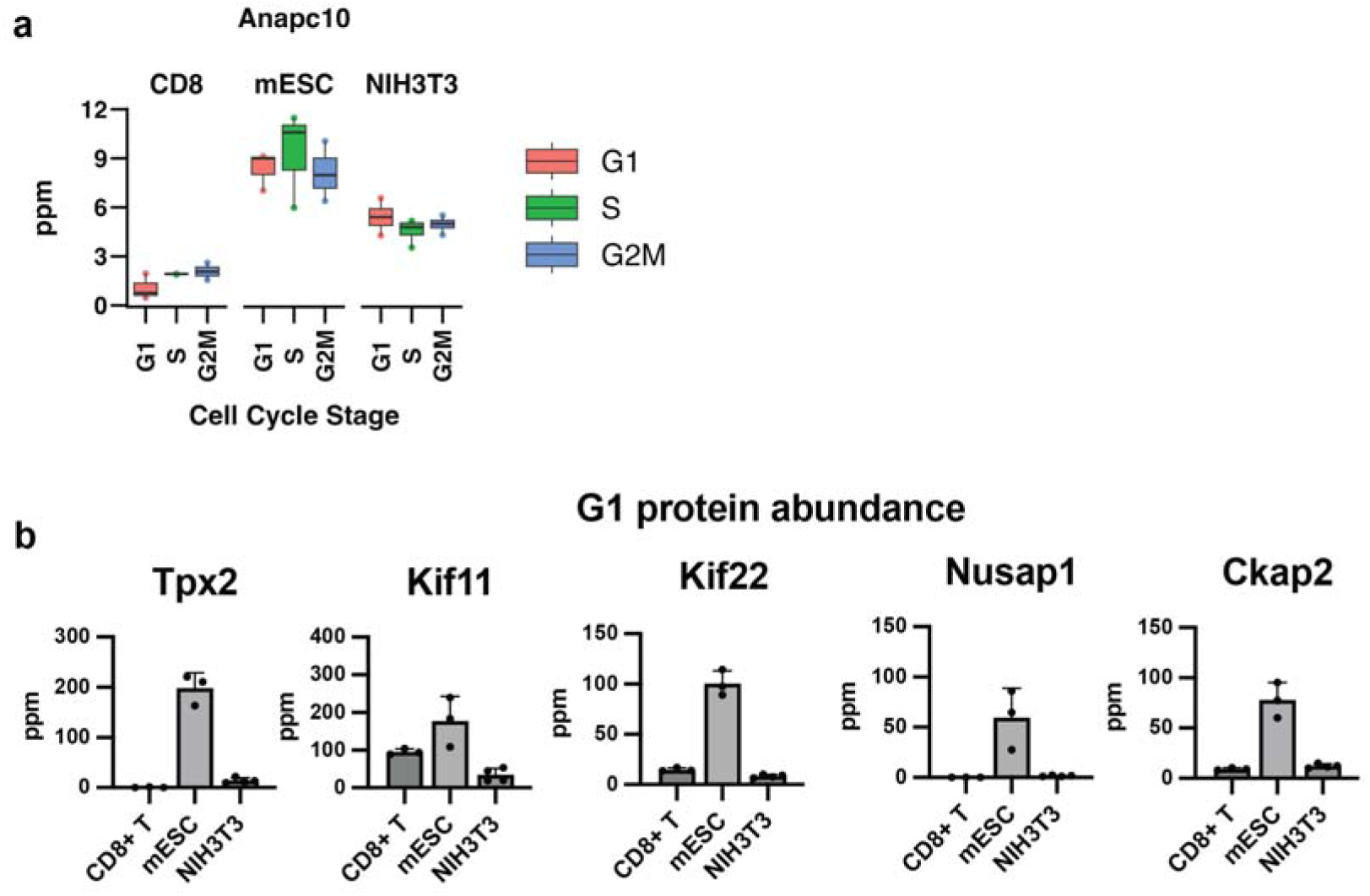
Comparing cell cycle regulation of the anaphase protein complex/cyclosome (APC/C) and its regulators. **a** Protein abundance (ppm) of Anapc10. **b** G1 phase protein abundance (ppm) of selected APC/C substrates. Error bars indicate standard deviation (n = 3-4).

**Supplementary Figure 2:**
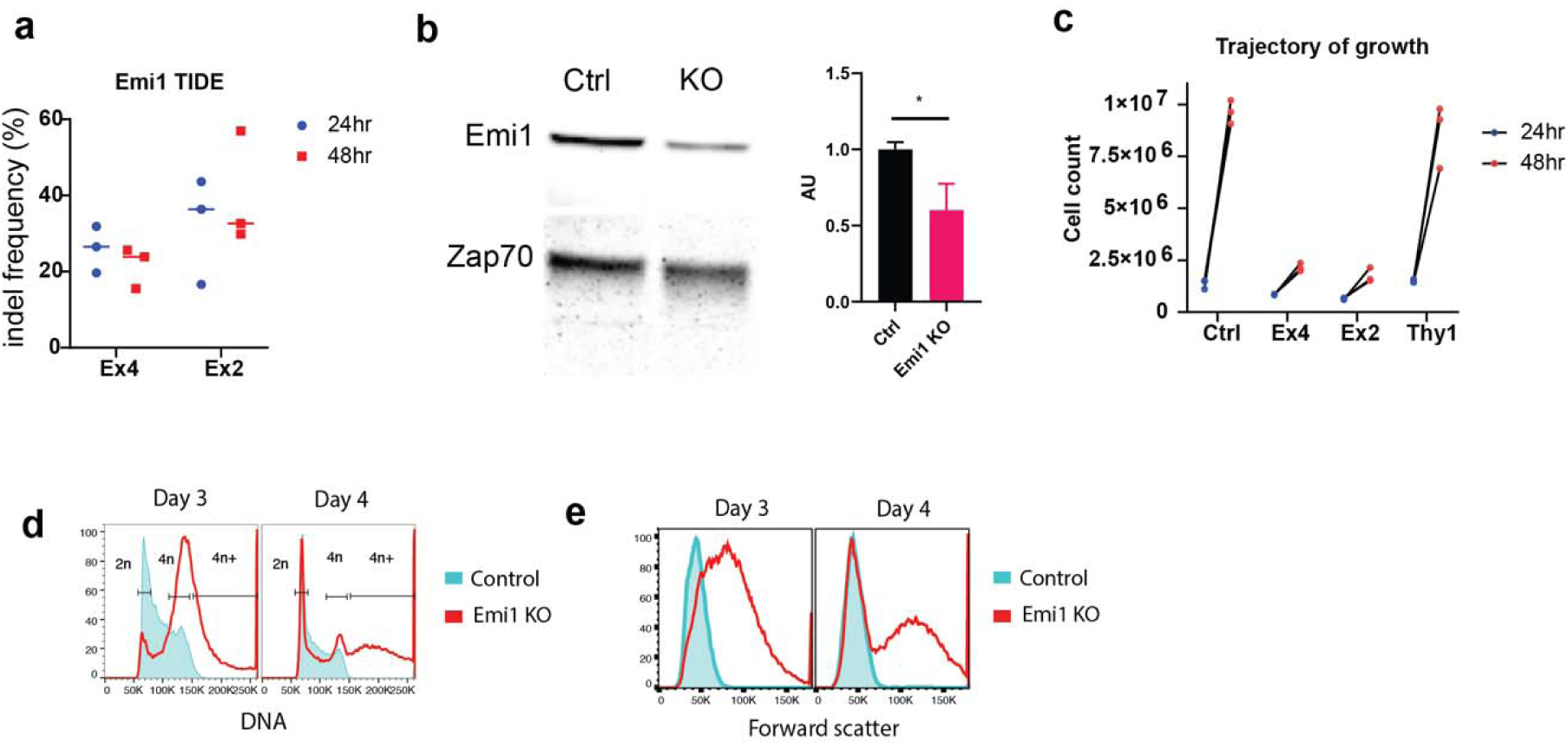
Emi1 promotes S-phase entry and safeguards against re-replication. **a** Emi1 knockout efficiency as measured by Tracking Indels by DEcomposition (TIDE) and PCR. Two sgRNAs were tested, which target exon 4 (Ex4) and exon 2 (Ex2) of the Emi1/Fbxo5 locus. **b** Immunoblot analysis of extracts from control and Emi1KO cells, probing for Emi1 protein. **c** Cell counts following control, Emi1KO, and Thy1KO. An sgRNA targeting Thy1 was used here as a negative control. Thy1 is a gene that expresses a cell surface marker that is not known to play a role in cell cycle progression in CD8 T cells. **d** DNA content histograms of control versus Emi1KO cells. **e** Forward scatter measurements of control and Emi1KO cells. Forward scatter is indicative of cell size.

**Supplementary Figure 3:**
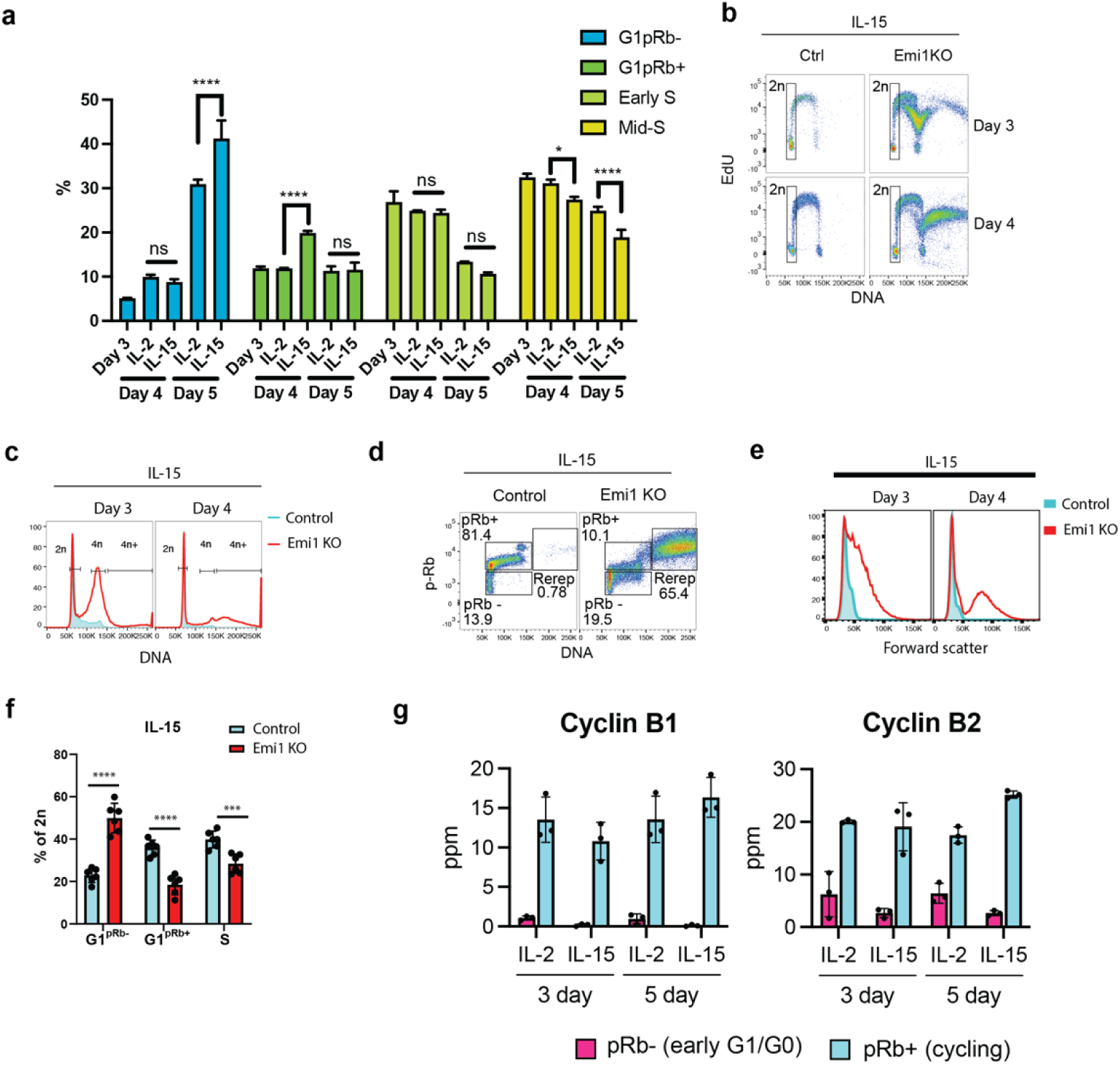
Role of Emi1 in memory-like CD8 T cells. **a** CD8 T cells were activated for two days and then cultured either in IL-2 or IL-15 for an additional 2 or 3 days. Cell cycle distribution was measured using phosphorylated Rb, EdU and DNA content. Total time post activation is shown on the x-axis. **b-f** OT-1 T cells were activated at day 0, transduced with the Cas9 complex with Emi1 targeted sgRNA or a blank control on day 2, cultured IL-15, and collected on Day 3 and Day 4 for analysis. EdU incorporation (b), DNA content histograms (c) Rb phosphorylation (d) forward scatter (e) and quantitation of early G1/G0 (G1^pRb-^), late G1 (G1^pRb+^), and early S-phase cells (f) are shown. Statistical significance was determined by unpaired Student’s t-test, NS = not significant, *p<0.05, **p<0.01, ***p<0.001, n = 6 mice from two independent cohorts (a-f), n = 3 mice (g). **g** Protein abundance (ppm) of Cyclin B1 and Cyclin B2 comparing IL-2 and IL-15 treated cells. (n = 3)

**Supplementary Figure 4:**
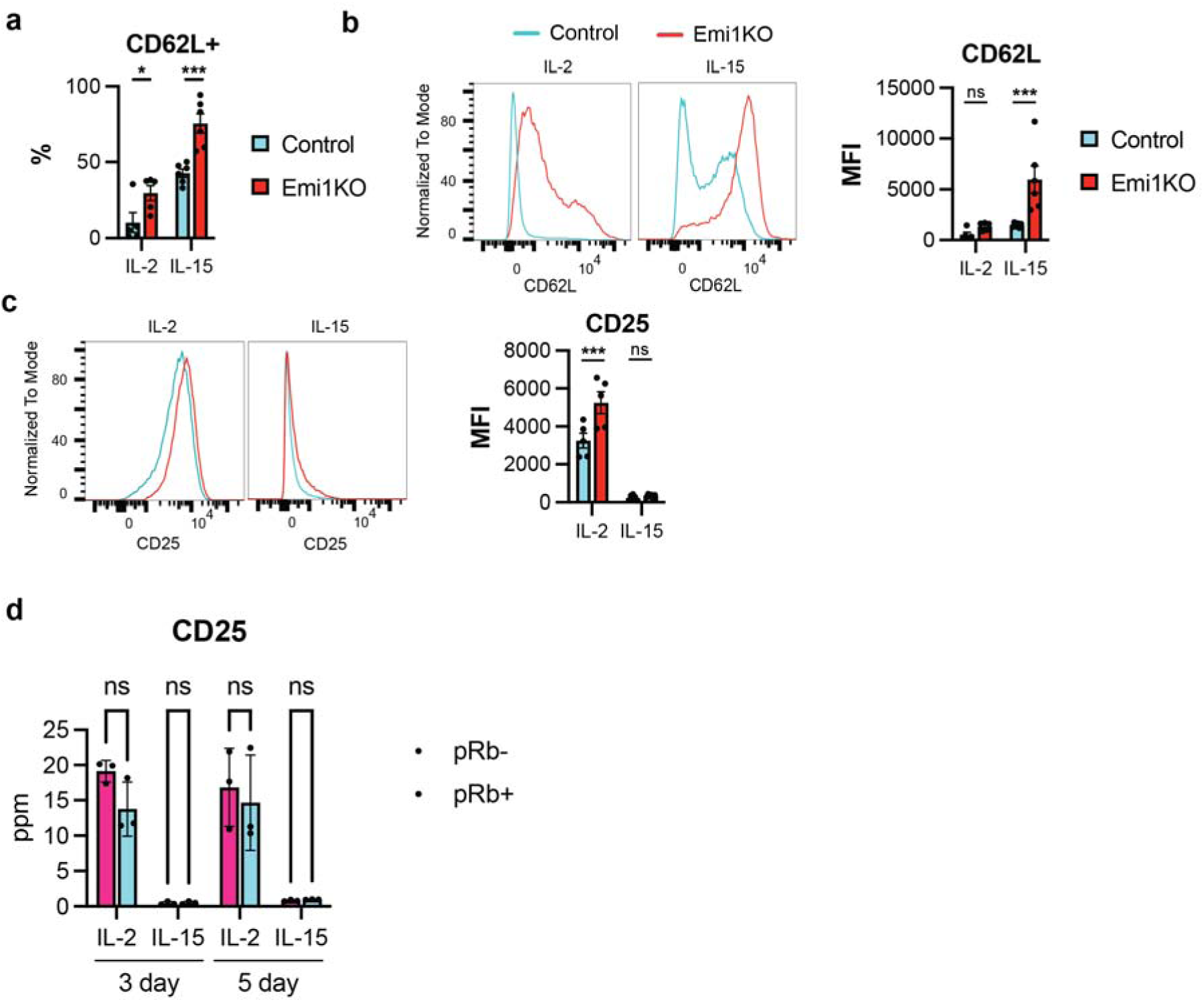
Emi1 controls T cell phenotype by regulating expression of CD62L. **a** CD8 T cells were activated, transduced with Cas9-sgRNA targeting Emi1 or a blank control on day 2, and then cultured in either IL-2 or IL-15 for an additional 2 days. (n = 5-6 animals across two independent animal cohorts). Quantitation of CD62L frequency. **b** Representative histograms of CD62L expression (left) and quantitation (right). **c** Representative histograms of CD25 expression (left) and quantitation (right). **d** Protein intensities (ppm) of CD25 in non-cycling (pRb-) and cycling (pRb+) CD8 T cells cultured in the indicate cytokines (IL-2 or IL-15). (n = 3). Statistical significance was determined by unpaired Student’s t-test, ns = not significant, *p<0.05, **p<0.01, ***p<0.001

